# Microtubule Inhibition Triggers MYC-Mediated Immunogenic Cell Death in Breast Cancer

**DOI:** 10.1101/2025.08.26.671755

**Authors:** Ruixian Liu, July Aung, Aino Peura, Antti O Hiltunen, Linda Id, Mariel Savelius, Martina Peltonen, Rita Turpin, Natasha Salmelin, Johanna M Anttila, Topi A Tervonen, Ilida Suleymanova, Daniel Nicorici, Minna Mutka, Johanna Mattson, Panu E. Kovanen, Laura Niinikoski, Tuomo Meretoja, Bina Prajapati, Andrei Goga, Pauliina M. Munne, Jeroen Pouwels, Juha Klefström

## Abstract

Oncogenic MYC promotes cancer cell proliferation, metabolism, and death, while also driving immunosuppression in the tumour microenvironment, complicating immune-based therapies. To counter MYC-driven immune evasion while leveraging MYC-dependent synthetic lethality (MYC-SL), we identified microtubule-targeting agents, including eribulin, as potent inducers of immunogenic cell death in MYC^high^ triple-negative breast cancer (TNBC). A screen of 528 oncology compounds using damage-associated molecular pattern (DAMP) reporters revealed that microtubule inhibitors induced key DAMPs, including HMGB1 secretion, calreticulin exposure, and double-stranded DNA release, leading to gasdermin-E associated cell death in MYC^high^ TNBCs. Immune cell co-culture assays showed immune activation, and patient-derived explant cultures confirmed pro-inflammatory cytokine responses. In vivo, cell-free media from eribulin-treated MYC^high^ murine TNBCs enhanced tumour protection in vaccination models compared to MYC-knockdown controls, linking MYC-dependent DAMP release to immunogenicity. These findings highlight a dual-function therapeutic strategy: agents that selectively induce MYC-dependent immunogenic cell death can provide both targeted cytotoxicity and local immune stimulation, thereby addressing a key limitation of conventional chemotherapeutics, offering a new approach for MYC-driven cancers.

## INTRODUCTION

MYC, a potent oncogenic transcription factor, is a master regulator of cellular growth, proliferation, and metabolism. Frequently overexpressed in cancers, MYC enforces cell cycle progression while orchestrating metabolic rewiring that fuels tumourigenesis^1–3^. However, despite its well-established role in promoting tumour growth, MYC harbours an inherent paradox: while driving proliferation, it simultaneously primes cells for apoptosis^4–6^. This duality presents a unique vulnerability–one that could be exploited to selectively eliminate MYC-driven tumours.

Cancer cells with high MYC expression develop specific dependencies to sustain their survival, including reliance on metabolic adaptations such as anaplerotic glutamine metabolism^2,7,8^. These dependencies expose vulnerabilities that can be exploited through MYC-dependent synthetic lethality (MYC-SL), a therapeutic framework focused on selectively eliminating MYC-driven tumour cells while minimizing toxicity to normal tissues^9,10^. By disrupting essential survival pathways, MYC-SL approaches amplify MYC’s inherent apoptotic sensitivity, leading to tumour-selective lethality. In fact, MYC-SL has been observed with metabolic inhibitors (CB-839, 2-deoxy-D-glucose), DNA damage response inhibitors (PARP, ATR inhibitors), cell cycle regulators (CDK, aurora kinase inhibitors), and BCL-2 inhibitors (venetoclax)^2,6,8,11,12^. Additionally, several chemotherapies, including microtubule-targeting agents (paclitaxel, vinblastine) and topoisomerase inhibitors (etoposide, doxorubicin) have been shown exhibiting MYC-SL effects^13–15^.

Beyond its role in proliferation and apoptosis, MYC profoundly influences tumour immunogenicity by shaping interactions within the tumour immune microenvironment (TIME) ^16,17^. MYC-driven breast cancers exhibit immune suppression through inhibition of the cGAS-STING pathway and downregulation of MHC-I expressin^18–21^. Additionally, MYC modulates chemokine networks and immune checkpoint proteins, fostering an immune-cold phenotype that enables tumour immune evasion^18,19,22^. This raises a compelling question: Could MYC’s pro-apoptotic tendencies be leveraged to reverse its immunosuppressive effects?

It is known that not all forms of cell death elicit the same immune response^23,24^. Immunogenic cell death (ICD), a specialized form of regulated stress-induced cell death, has the potential to transform an immune-cold tumour into an immune-hot one. ICD triggers the release of tumour antigens and damage-associated molecular patterns (DAMPs), including high mobility group box 1 (HMGB1) and calreticulin (CALR), both of which activate dendritic cells (DCs) and enhance antigen presentation^25,26^. These inflammatory signals initiate robust anti-tumour immunity, offering a promising avenue to enhance cancer immunotherapy^27^. Given MYC’s role in apoptosis and immune modulation, we sought to determine whether MYC-SL mechanisms could also induce ICD – effectively flipping MYC-driven immunosuppression into immunoactivation.

Rather than first identifying MYC-SL chemotherapies and subsequently assessing their immunogenic potential, here we prioritized an unbiased screen for ICD inducers. This approach allowed us to identify compounds capable of triggering robust immune activation, independent of preconceived assumptions about MYC-SL. By screening a library of 528 approved and investigational oncology drugs for DAMP exposure, we identified microtubule-targeting agents as potent ICD inducers. Candidates were validated in vitro by assessing their ability to activate antigen-presenting and T-cell activating HLA-DR⁺ CD86⁺ DCs. Our findings revealed that microtubule-targeting agents significantly increased activated DC populations in co-culture, with indications of gasdermin E-mediated pyroptosis as a potential underlying mechanism. Finally, using breast cancer patient-derived explant cultures (PDECs) and tumour vaccination mouse models, we demonstrate that MYC-driven tumours exhibit enhanced immunogenicity in response to microtubule-targeting therapy – suggesting that MYC-dependent cell death could indeed be harnessed to switch immune suppression into immune activation.

## RESULTS

### Oncology Drug Screen Identifies Compounds inducing Damage-Associated Molecular Pattern (DAMP) exposure in TNBC cells

To identify small-molecule compounds capable of inducing damage-associated molecular pattern (DAMP) exposure in triple-negative breast cancer (TNBC) cell lines, we screened a curated library of 528 approved and emerging investigational oncology drugs **(Table S1)** using DAMP-reporter TNBC models. A dual-reporter system was established by fusing HMGB1 to green fluorescent protein (GFP) and calreticulin (CALR) to red fluorescent protein (RFP), enabling real-time visualization of DAMP dynamics^28,29^. These reporters were stably expressed in two TNBC cell lines, HCC38 and MDA-MB-231, selected for their flat morphology and suitability for high-resolution imaging **(Fig. 1a)**. Mitoxantrone (MTX), a type II topoisomerase inhibitor and well-characterized ICD inducer^25,30^, served as positive control to validate DAMP signal induction and assay performance **(Fig. 1b)**. Quantification of HMGB1 nuclear depletion and CALR membrane translocation relative to controls confirmed robust MTX-induced DAMP exposure **(Figs. 1c, d)**.

**Figure 1.**
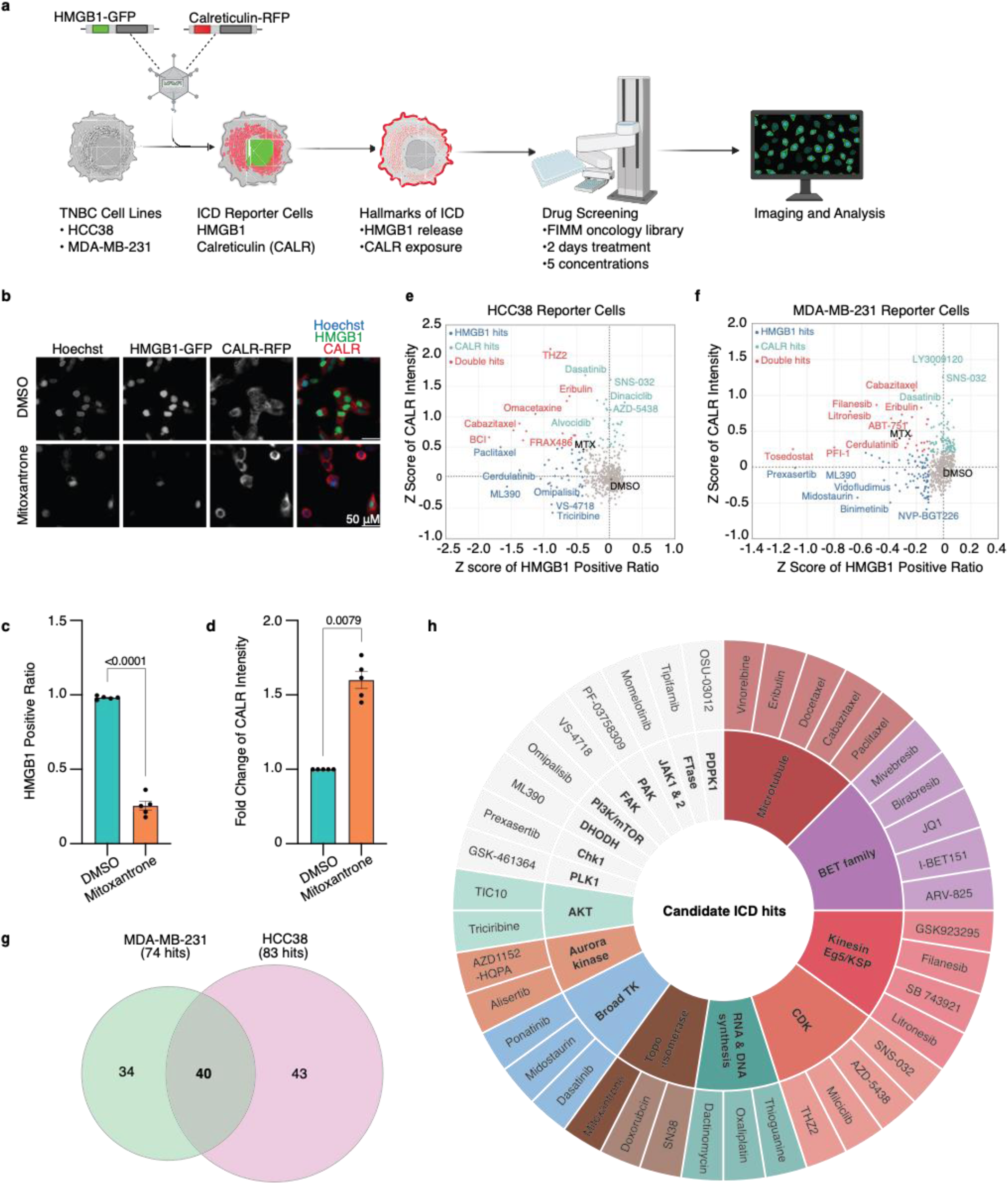
Imaging-based screen identifies candidate ICD-inducing compounds in TNBC cells. **a.** Schematic workflow of the high-content imaging screen using a library of approved and investigational oncology compounds in dual DAMP-reporter TNBC cell lines (HCC38 and MDA-MB-231). **b.** Representative images of HCC38 reporter cells treated for 48 hours with 2.5 µM Mitoxantrone or 0.1% DMSO and 4% PFA fixation (vehicle control). Scale bar = 50 µm. **c-d.** Quantification of (c) HMGB1 nuclear depletion (% HMGB1-positive cells; Welch’s T-test) and (d) CALR membrane translocation (fold-change in CALR intensity; Mann-Whitney U-test) after 48 hours treatment with Mitoxantrone or DMSO. Error bars represent SEM. **e-f.** Z-score comparisons of (e) HMGB1-positive cell ratio and (f) CALR intensity in HCC38 and MDA-MB-231 reporter cell lines. Top 10% of compounds for HMGB1 and CALR are highlighted in blue and green, respectively; compounds scoring in the top 10% for both metrics are shown in red. **g.** Venn diagram illustrating the overlap of candidate compounds identified in both cell lines. **h.** Sunburst plot classifying top-ranked ICD candidate compounds by their putative molecular targets.

Each compound was tested at five concentrations ranging from 1 nM to 10 μM over 48 hours, and DAMP responses were evaluated by high-content imaging. DAMP induction was defined as nuclear depletion of HMGB1-GFP and plasma membrane translocation of CALR-RFP. Fluorescent intensities were normalized to DMSO control values and converted to Z-scores (z = (x – μ)/σ) to quantify DAMP induction across treatments **(Figs. 1e, f)**. Ranked by Z-score, the top 40 compounds demonstrating consistent DAMP responses across both cell lines were selected for downstream validation **(Figs. 1g, h)**. The image-based profiling revealed distinct clusters (n ≥ 3) of DAMP-inducing compounds across diverse drug classes, including microtubule inhibitors, BET protein modulators, kinesin and kinesin spindle protein (KSP) inhibitors, cyclin-dependent kinase (CDK) inhibitors, along with agents targeting nucleic acid synthesis, topoisomerases, and tyrosine kinases **(Fig. 1h)**.

### Microtubule-Targeting Compounds Promote Dendritic Cell Maturation via Immunogenic Cell Death

DAMPs released during ICD activate pattern recognition receptors, such as toll-like receptors, on DCs, promoting their maturation^28^. DC maturation is characterized by reduced phagocytic activity, upregulation of MHC class I and class II and costimulatory molecules (CD40, CD54, CD80, CD86), and increased secretion of pro-inflammatory cytokines, including IL-18 and TNF-α – features essential for priming antigen-specific T-cell responses^31^. To assess whether the top 40 compounds identified in **Fig. 1h** induce functional DC activation, we performed immunophenotyping and phagocytosis assays. CD14^+^ monocytes were isolated from peripheral blood mononuclear cells (PBMCs) and differentiated into immature monocyte-derived DCs (moDCs). MoDCs were either co-cultured with carboxyfluorescein succinimidyl ester (CFSE)-labelled TNBC cells pre-treated with candidate compounds for 24 hours or exposed to conditioned medium (CM) collected from treated TNBC cultures **(Fig. 2a)**. DC maturation was assessed by flow cytometry based on surface expression of activation markers CD86 and HLA-DR, while phagocytic activity was measured by the uptake of CFSE-labelled tumour cells.

**Figure 2.**
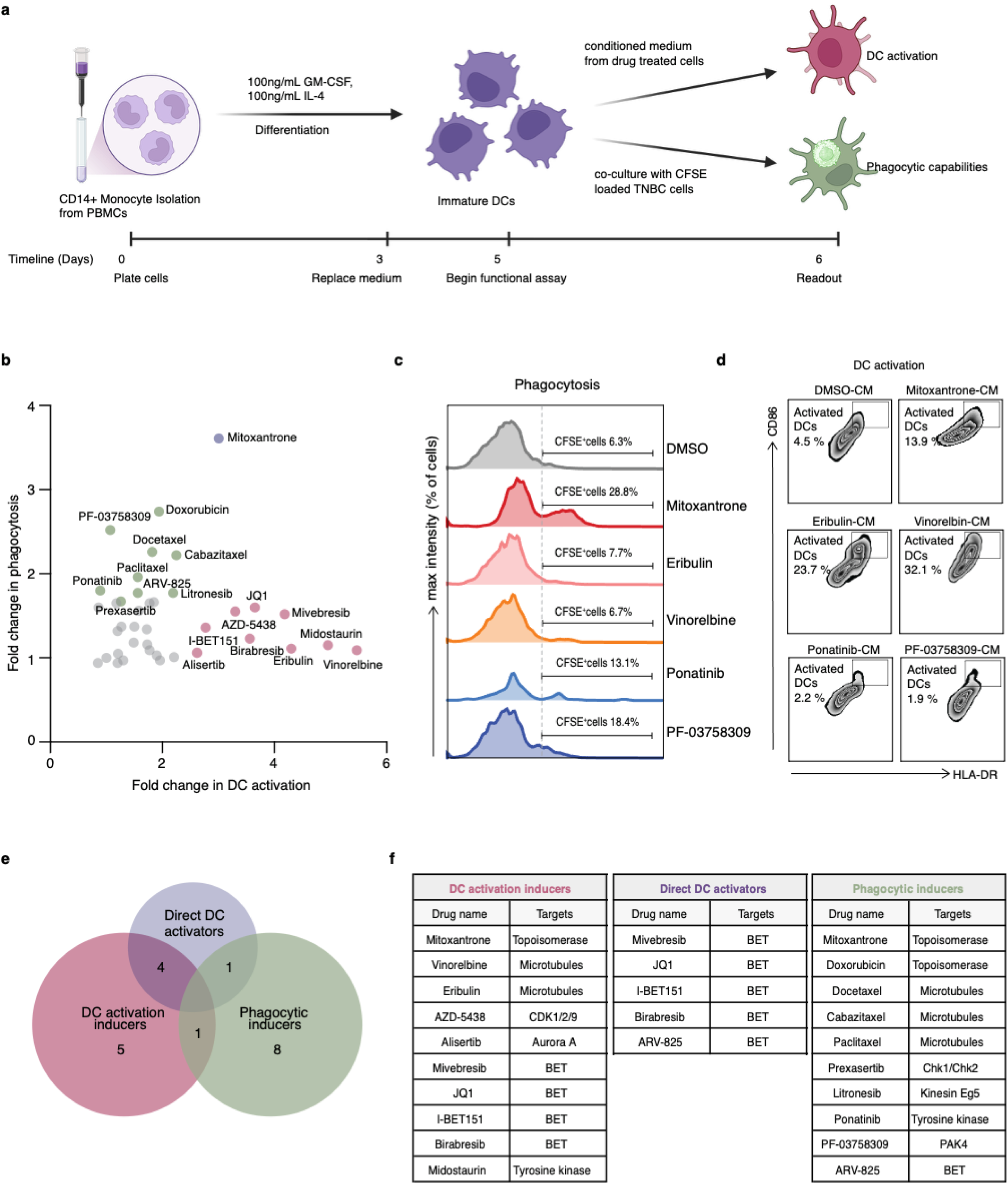
Dendritic cell-based functional validation unveils enrichment of ICD targets linked to mitosis. **a.** Schematic workflow representing dendritic cell (DC) isolation from PBMC-derived monocytes and DC functional assay with either conditioned medium or drug-treated CFSE^+^ tumour cells. **b.** Fold change in phagocytic and DC activation markers for 40 candidate ICD hits. Top 10 hits for phagocytosis marked in green, DC activation marked in red, and dual inducer, Mitoxantrone, marked in purple. **c-d.** Flow cytometric analysis of fold change in CFSE expression for phagocytosis and DC activation markers following functional DC assays. **e-f.** Compounds categorized according to their activation profiles.

Live cell imaging revealed a progressive increase in immunogenic activity beginning approximately 24 hours post-treatment **(Fig. 4a).** These findings suggest that DC maturation begins around 20-24 hours after actively taking up immunogenic material from dying tumour cells. This timing aligns with previous reports showing a phased pattern of phagocytosis and maturation kinetics that vary depending on the type of DC and the nature of stimuli involved^32,33^. Compounds were therefore ranked based on combined fold-changes in DC activation and phagocytic activity to identify the most potent ICD inducers, acknowledging that full DC maturation may not have been achieved at the experimental endpoint **(Fig. 2b)**.

Mature DCs capable of antigen presentation and T-cell priming are critical for initiating adaptive immune responses. Among tested compounds, MTX-treated TNBC cells elicited the strongest phagocytic response, with >3-fold increase in CFSE^+^ DCs compared to eribulin (ERI)-, or vinorelbine (VINO)-treated cells **(Fig. 2c)**. In contrast, when DCs were exposed to drug-CM, VINO-CM most effectively induced DC maturation, increasing the proportion of CD86^+^ HLA-DR^+^ cells by 8-fold (45.1%) relative to vehicle control (DMSO-CM). Comparatively, ERI-CM and MTX-CM induced more modest increases of 28.7% and 20.6%, respectively **(Fig. 2d)**.

The top 10% of compounds were further classified based on their DC activation profiles. To exclude confounding effects of direct drug toxicity, immature DCs were also exposed to the compounds in the absence of tumour cells. Flow cytometric profiling revealed that several BET inhibitors – compounds with known antineoplastic activity, directly activated DCs independent of tumour-derived signals. These were therefore classified separately as “direct DC activators” **(Figs. 2e, f)**. Microtubule inhibitors emerged as the most effective drug class, representing 25% of the top-performing compounds and reinforcing the link between mitotic disruption and ICD by proxy of DAMPs release. While most compounds enhanced either phagocytosis or DC activation, MTX notably displayed a strong proclivity for both, underscoring its potency as an ICD inducer. These findings highlight the duality of mitotic regulators – not only as inducers of tumour-derived DAMPs, but also as modulators of innate immune activation – reinforcing their central and emerging role in ICD.

### Microtubule Inhibition Induces MYC-directed Synthetic Lethal Immunogenic Cell Death

MYC accelerates mitotic progression, increasing cellular dependence on precise spindle function. This heightened reliance creates a specific vulnerability to microtubule-targeting agents such as taxanes^13,34^. This drug class – including paclitaxel, carbazitaxel, and docetaxel – has been a cornerstone of chemotherapeutic regimens since the 1990s and remains widely used in first-line settings^35,36^. Nonetheless, their clinical efficacy remains constrained by the frequent development of drug-resistance and by dose-limiting toxicities. To address these limitations, we investigated whether the top ICD compounds also exhibited MYC-synthetic lethality (MYC-SL), thereby selectively targeting MYC^high^ cancer cells and enabling the identification of agents with potentially improved safety profiles.

Using the Cancer Cell Line Encyclopedia (CCLE), we stratified pan-cancer cell lines into MYC^high^ and MYC^low^ subtypes based on the “MYC V1V2” signature^37^ which integrates the Hallmark gene sets, “HALLMARK_MYC_TARGETS_V1” and “HALLMARK_MYC_TARGETS_V2”^38^. The signature correlates strongly with elevated MYC protein levels, making it a robust surrogate marker for MYC activity. We then queried drug sensitivity data from the 24Q2 PRISM Repurposing Screen–where compounds were tested at 2.5 μM for five days and viability inferred from mRNA barcode abundance relative to DMSO controls ^39^ to assess the performance of our lead ICD inducers. Sensitivity profiles were obtained for 16 top-scoring ICD inducers across 146 MYC^high^ and 162 MYC^low^ cell lines ranked by MYC activity. Of these, 13 compounds significantly reduced viability preferentially in MYC^high^ cell lines, congruent with MYC synthetic lethality **(Fig. 3a)**. Notably, microtubule inhibitors emerged as the most consistent drivers of MYC^high^ selective cytotoxicity in the PRISM dataset **(Fig. 3b)**. To validate the results, we performed independent cell viability assays using CellTiter-Glo across three MYC^high^ and three MYC^low^ TNBC cell lines^37^. Drug sensitivity scores, calculated as previously described^37,40^, confirmed significantly greater sensitivity in MYC^high^ cell lines **(Fig. 3c)**, consistent with PRISM predictions and supporting the MYC-SL activity of microtubule-targeting agents.

**Figure 3.**
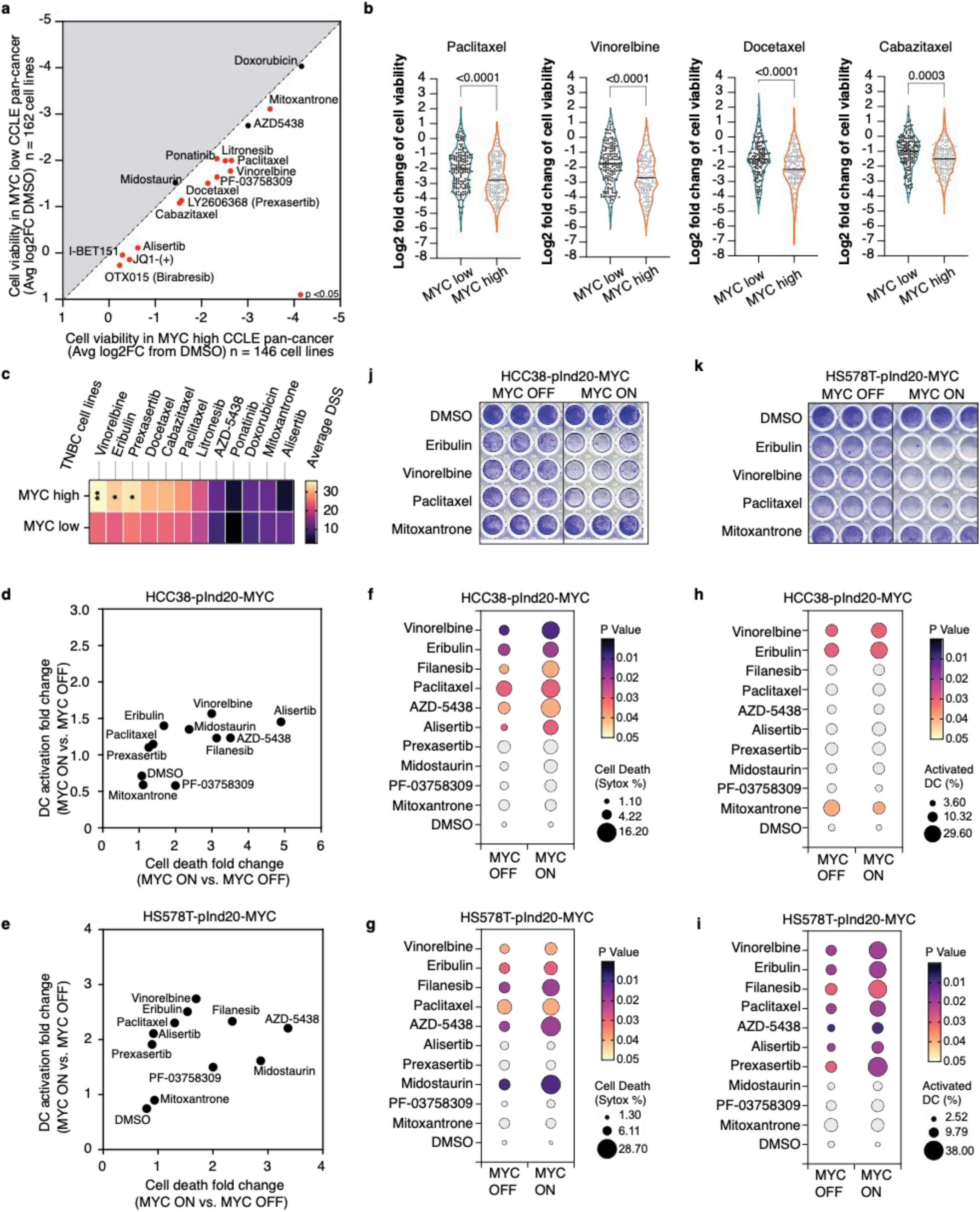
Microtubule-targeting agents induce MYC-directed ICD. **a.** Average Log2-fold change in cell viability (DMSO vs. treatment) in MYC^high^ (n = 146) and MYC^low^ (n = 162) CCLE pan-cancer cell lines ranked by MYC V1V2 signature score. MYC-SL compounds show significantly lower viability in MYC^high^ cell lines compared to MYC^low^ cell lines (Red, Student’s T-test, P < 0.05). **b.** Comparison of cell viability between MYC^high^ (n = 146) and MYC^low^ (n = 162) CCLE pan-cancer cell lines treated with four microtubule-targeting agents. **c.** Drug sensitivity score (DSS) comparison between MYC^high^ (HCC1395, HCC1806, and BT-549) and MYC^low^ (HCC1143, MDA-MB-231, and MDA-MB-468) TNBC cell lines for top ICD candidate drugs (*P < 0.05, **P < 0.01, Multiple unpaired T-tests). **d-e.** Fold-changes in cell death and DC activation with or without doxycycline induction of MYC in HCC38-pInd20-MYC (d) and Hs578T-pInd20-MYC (e) cell lines. **f-g.** Comparison of cell death (Sytox green %) between MYC ON (+ 100 nM Doxycycline) and MYC OFF (PBS) conditions in HCC38-pInd20-MYC (f) and Hs578T-pInd20-MYC (g) cell lines treated with top ICD candidate compounds. **h-i.** Comparison of DC activation (HLA-DR+ CD86+) in HCC38-pInd20-MYC (h) and Hs578T-pInd20-MYC (i) cell lines after treatment with top ICD candidate compounds under MYC ON and OFF conditions. **j-k.** Cell viability assessed by Coomassie blue staining in HCC38-pInd20-MYC (j) and Hs578T-pInd20MYC (k) cell lines with differential MYC induction.

To directly assess the impact of MYC expression on both cell death and ICD, we employed complementary assays. Cell death was measured using SYTOX Green, a membrane-impermeant nucleic acid dye that selectively stains cells with compromised plasma membranes, and ICD by presence of dendritic cell (DC) activation markers. We engineered two triple-negative breast cancer (TNBC) cell lines, HCC38 and Hs578T, to express doxycycline-inducible MYC. These cells were treated with the top 10 candidate compounds, and after 48 hours, both SYTOX Green fluorescence and DC activation were quantified using flow cytometry. Fold-changes between MYC^high^ and MYC^low^ conditions were then compared **(Figs. 3d, e)**. Across both assays and in both cell models, microtubule-targeting compounds induced greater immunogenic cytotoxicity in MYC-induced cells relative to controls, as reflected by increased SYTOX Green staining and elevated DC activation marker expression. On average, MYC induction doubled cytotoxic responses in both models **(Figs. 3f–i)**. Furthermore, Coomassie staining of adherent cells 2-days post-treatment revealed a 45–60% decrease in viable cells in the MYC ON condition, in striking contrast to the MYC OFF condition, where no such effect was observed **(Figs. 3j, k)**.

Collectively, our findings show that microtubule inhibitors selectively target MYC^high^ cancer cells, doubling cytotoxic and immune-activating responses upon MYC induction, thereby linking MYC overexpression to heightened vulnerability to microtubule disruption and enhanced ICD.

### Microtubule Inhibitors Induce Gasdermin E Cleavage and Lytic Immunogenic Cell Death

Live-cell imaging of HCC38 ICD-reporter cells treated with eribulin revealed striking morphological changes ∼24 h post-treatment. Initially flat, adherent cells rounded and enlarged, developing persistent bubble-like protrusions accompanied by progressive loss of nuclear HMGB1-GFP and translocation of CALR-RFP (**Fig. 4a**). After several hours of swelling, cells underwent abrupt membrane rupture, leaving membrane fragments that degraded slowly. This lytic sequence contrasted with classical apoptosis, which features discrete membrane blebbing and apoptotic body formation.

**Figure 4.**
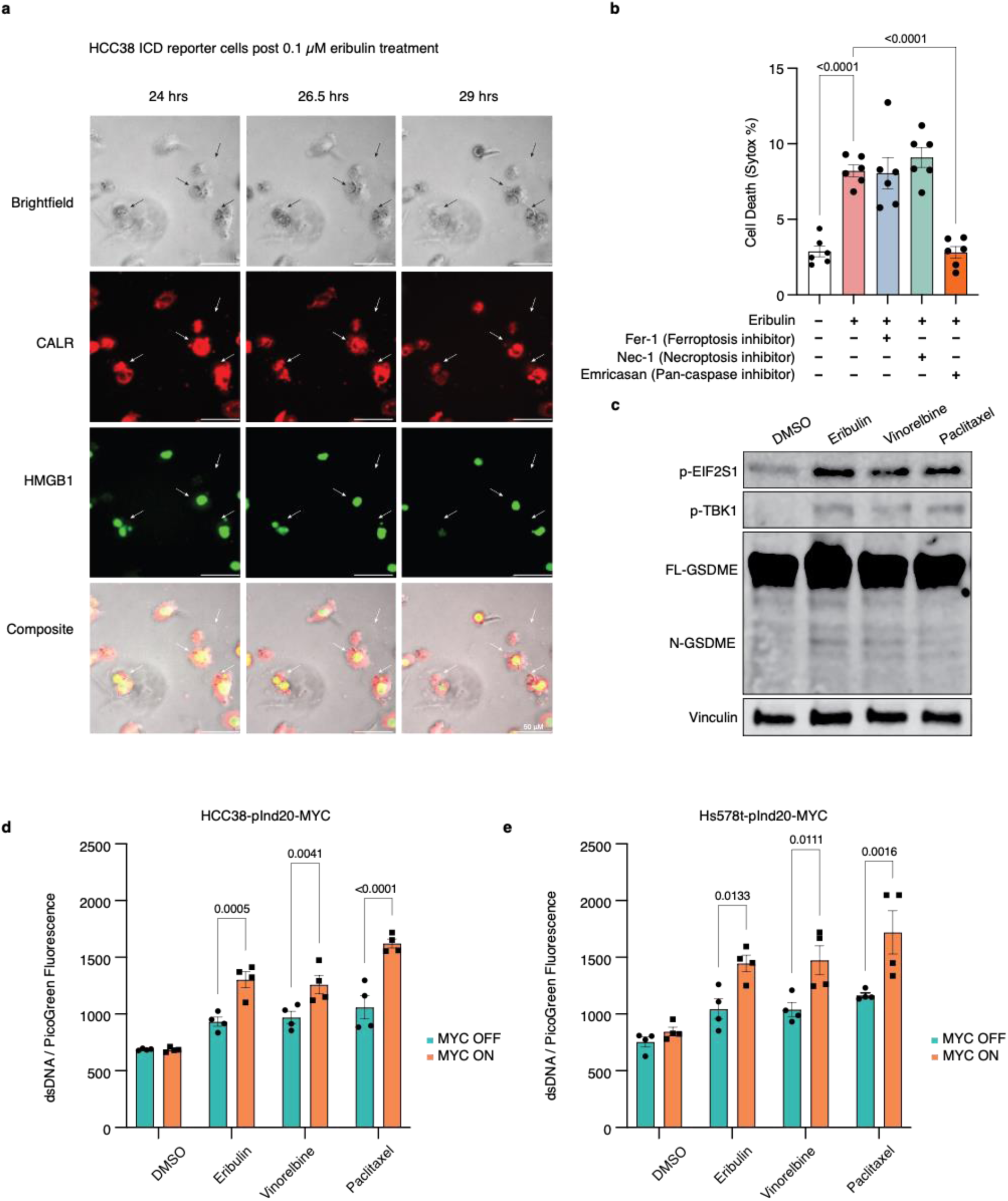
Microtubule-targeting treatments induce Gasdermin-E cleavage and pyroptosis. **a.** Morphological and phenotypic changes observed in HCC38 ICD reporter cells post 0.1 µM eribulin treatment under time-lapse microscopy, scale bar = 50 µM. Arrows indicate swelling cells accompanied by loss of HMGB1 and CALR signals. **b.** Percentage of SytoxGreen-positive Hs578T cells following pre-treatment with various cell death inhibitors and subsequent treatment with 0.1 µM Eribulin, error bars represent SEM. **c.** Representative Western blot showing Gasdermin E cleavage after 24 hours of treatment with 0.1 µM Eribulin, 0.1 µM Vinorelbine, or 0.1 µM Paclitaxel in Hs578T. **d-e.** dsDNA concentration in the supernatant, measured with PicoGreen, from HCC38-doxMyc and Hs578T-doxMyc cells treated with 0.1 µM Eribulin, 0.1 µM Vinorelbine, or 0.1 µM Paclitaxel for 24 hours. P-values calculated using 2-way ANOVA, followed by Holm-Sidak’s multiple comparisons test, error bars represent SEM.

In pharmacological rescue assays with Hs578T cells, inhibition of necroptosis (Nec-1) or ferroptosis (Fer-1) did not block eribulin-induced death. In contrast, the pan-caspase inhibitor emricasan significantly improved viability (p < 0.0001) (**Fig. 4b**), pointing to a caspase-dependent mechanism.

Western blotting detected cleavage of gasdermin E (GSDME) and release of its pore-forming N-terminal fragment following treatment with multiple microtubule inhibitors, including eribulin (**Fig. 4c).** Although N-GSDME was present at low levels, phosphorylated eIF2α (p-EIF2S1) and TBK1 (p-TBK1)–markers of integrated stress and inflammatory signalling–were also detected. To assess membrane rupture more directly, we quantified extracellular double-stranded DNA, one hallmark of lytic pyroptosis^41,42^. Across eribulin, vinorelbine, and paclitaxel treatments, HCC38 and Hs578T cells with doxycycline-induced MYC released significantly more double stranded DNA (dsDNA) than non-induced controls (**Figs. 4d, e**).

Taken together, the observed swelling, explosive rupture, caspase dependence, GSDME cleavage, and dsDNA release are most congruent with a lytic form of ICD that bears the morphological and molecular hallmarks of GSDME-mediated pyroptosis.

### Eribulin Elicits Immunogenic Cytokine Signatures in MYC^high^ Breast Cancer Explants with Physiologic Tumour Immune Microenvironment

To validate our in vitro findings, we tested ICD-inducing agents in three-dimensional (3D) ex-vivo Patient-Derived Explant Cultures (PDECs) **(Fig. 5a)**. PDECs are tumour fragments cultured directly from patient tissue, preserving the tumour’s native architecture and local immune microenvironment, including immune cell composition and signalling networks. Both recent studies from our group and others have shown that short-term PDECs maintain immune cell infiltration, cytokine profiles, and tumour heterogeneity, providing an accurate model to study tumour biology, immune responses, and the efficacy of therapies, including immunotherapies and combination treatments^43–46^.

**Figure 5.**
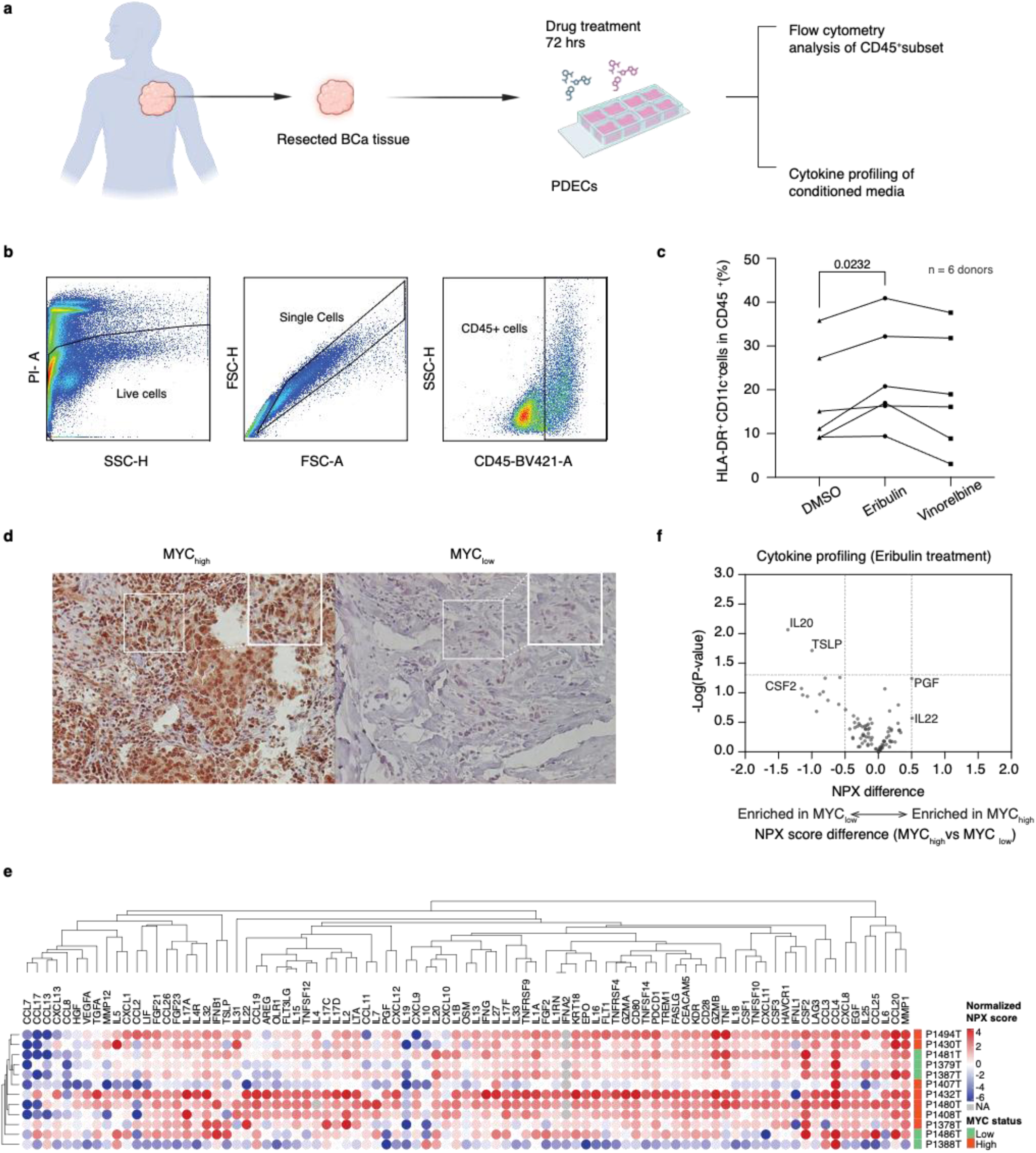
Eribulin treatment induces a robust immunogenic signature in MYC-high patient-derived explants. **a.** Schematic workflow of patient-derived explant cultures (PDECs) following 3 days drug treatment. **b.** Flow cytometry gating for identifying dendritic cell subsets in 6 PDECs. **c.** Graph indicates proportion of dendritic cell subsets in 6 donor explants after different drug treatments. **d.** Representative IHC images of FFPE MYC^high^ and MYC^low^ breast tumours stained for MYC (Abcam Y69 antibody). Inset highlights nuclear MYC signal. **e.** Cytokine and chemokine expression profile in 7 MYC^high^ and 5 MYC^low^ PDECs treated with 1 µM Eribulin. NPX stands for normalized protein expression. Hierarchical clustering conducted on both cytokines and patient groups by One minus Spearman rank correlation analysis, based on average linkage method. Heatmap generated with Morpheus, https://software.broadinstitute.org/morpheus. **f.** Volcano plot showing differential cytokine expression in MYC^high^ versus MYC^low^ PDECs following 1 µM Eribulin treatment.

Flow cytometric analysis of tumour-infiltrating immune cells (CD45^+^ leukocytes) **(Figs. 5b, c)** revealed that treatment with eribulin consistently increased the proportion of CD11c^+^ DCs expressing HLA-DR⁺, a key MHC-II molecule involved in antigen presentation to CD4^+^ T helper cells. This upregulation of HLA-DR⁺ was observed in 5 out of 6 PDEC samples, indicating a robust DC activation in response to eribulin. In contrast, vinorelbine treatment produced a more variable effect on HLA-DR⁺ expression on DCs across these samples **(Fig. 5c)**. Elevated HLA-DR expression on CD11c⁺ DCs demonstrates enhanced antigen presenting capacity required for promoting downstream adaptive immune responses, potentially improving anti-tumour immunity following eribulin therapy.

Next, we collected a new series of 12 PDECs, stratifying samples as MYC^high^ or MYC^low^ based on immunohistochemical scoring of nuclear MYC intensity and the proportion of positive cells in the original tumour tissue **(Figs. 5d, S1)**. To investigate eribulin-induced local inflammatory responses, conditioned media from seven MYC^high^ and five MYC^low^ PDECs treated with 1 μM eribulin were analysed using Olink’s immune-oncology cytokine panel **(Figs. 5e, f)**. Cytokine and chemokine profiling revealed that eribulin activated broad secretion of immunomodulatory cytokines in PDECs **(Fig. 5e)**. Notably, we observed lower expression of cytokines associated with immunosuppressive and regulatory functions, including IL-19 and IL-10, which are linked to anti-inflammatory signalling and T-cell suppression. In addition, lower expression characterised CCL13 and CCL17, which are associated with recruitment of M2-like macrophages and pro-tumoral immune evasion ^47–50^. In contrast, several inflammatory and cytotoxic effector cytokines were strongly activated, including GZMB, TNF, IL-6, CCL4, and CXCL8 – key mediators of pro-inflammatory immune activation. These cytokines are known to promote recruitment and activation of DCs, cytotoxic T-cells and natural killer (NK) cells central to anti-tumour immunity^51,52^. Together, these findings suggest that eribulin treatment remodels the cytokine milieu of PDECs to promote immune activation and potentially enhance anti-tumour immunity.

Moreover, in MYC^high^ PDECs, PGF and IL-22 were upregulated, whereas CSF2, IL-20, and TSLP were downregulated compared to MYC^low^ PDECs. The upregulation of PGF (PDGF) and IL-22 suggests enhanced tumour survival and angiogenic signalling^53,54^, while downregulation of CSF2 (GMCSF), IL-20, and TSLP may reflect reduced immune-activating signals^55–57^, consistent with the recently documented immunosuppressive role of MYC^18,19^. Despite these differences, the overall cytokine profiles between MYC^high^ and MYC^low^ PDECs remained largely similar, indicating that eribulin-induced ICD still occurs in the PDEC model, with local immune-activating effects, even in the presence of MYC expression. However, except for IL-20 and TSLP, the observed differences were not statistically powered possibly due to a small sample size, and future studies are needed to clarify these trends.

### MYC Is Essential for Eribulin-Induced Immunogenic Cell Death and Antitumour Immunity In Vivo

To assess whether the eribulin-induced, MYC-dependent synthetic lethal ICD death observed in vitro elicits functional antitumour immunity in vivo, we developed an in vivo prophylactic vaccine model in immunocompetent Balb/cJRJ mice. Prior attempts using lysate-based whole-cell vaccines derived from dying tumour cells consistently led to tumour formation at the vaccine site during post-vaccine recovery, likely due to residual viable cells (**Fig. S2a**), which precluded subsequent tumour challenge experiments. To circumvent this, we generated a cell-free vaccine enriched with immunogenic DAMPs derived from dying 4T1 tumour cells treated with microtubule-targeting agents.

To evaluate the impact of MYC on ICD vaccine efficacy, 4T1 cells which endogenously express high levels of MYC^58^ were transduced with pLKO.1-shMYC, generating 4T1-pLKO.1-shMYC cells with stable MYC silencing **(Figs. 6a, b)**. MYC^high^ and MYC-knockdown cells were treated with eribulin for 24 hours, after which conditioned medium was harvested and concentrated up to 100-fold through multiple ultrafiltration and ultracentrifugation steps. Using Amicon^®^ ultrafilters with varying molecular weight cutoffs, we isolated a concentrated, fractionated, cell-free conditioned medium (FCM), enriched in key immunogenic DAMPs including HMGB1, and CALR – hereafter referred to as FCM-vaccine (FCM-vac) **(Fig. 6c)**.

**Figure 6.**
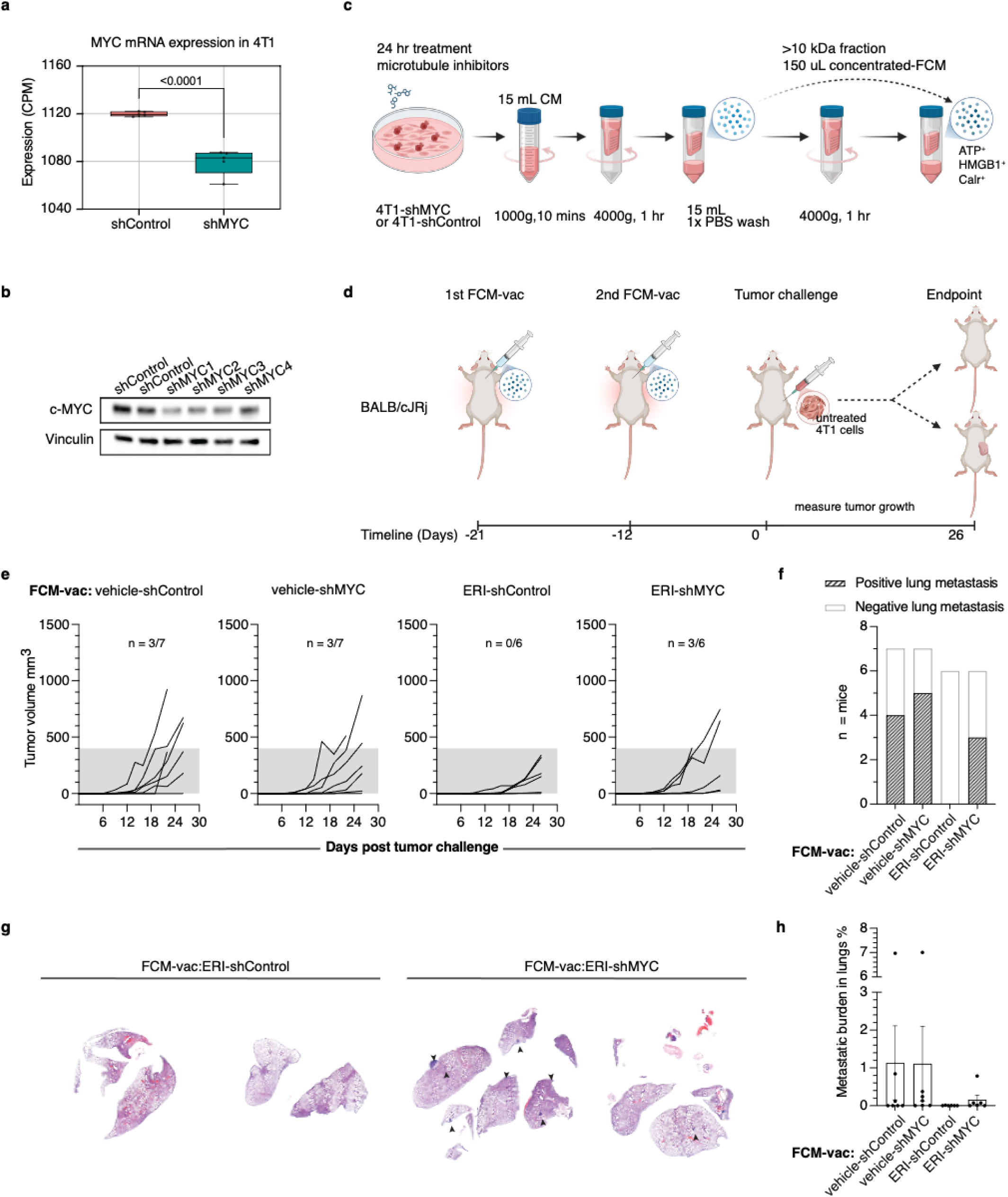
MYC expression is essential for Synthetic Lethal Immunogenic Cell Death in vivo. **a.** MYC mRNA expression by counts per million in 4T1 cells before and after shRNA silencing. Student’s T-test. **b.** Corresponding western blot analysis of MYC protein levels after silencing with different shRNA sequences in 4T1 cell lines, shMYC2 construct was selected for in vivo experiments. **c.** Schematic overview of experimental workflow generating concentrated, fractionated, cell free conditioned medium (FCM) for vaccination assays. **d.** Graphical representation of vaccine dosing schedule and in vivo tumour challenge assays in BALB/cJRj mice. **e.** Tumour growth curves in BALB/cJRj mice vaccinated with FCM derived from Eribulin or vehicle (DMSO) treated 4T1-pLKO.1-shMYC and 4T1-pLKO.1-shControl cells. N = number of mice with tumours >400mm^3^. **f.** Incidence of breast cancer metastasis to lung tissue in mice of DMSO and Eribulin treated FCM-vac groups. **g.** Representative H&E staining of mice lung tissues after vaccination with FCM of Eribulin-treated 4T1-pLKO.1-shMYC, and 4T1-pLKO.1-shControl cells. Arrows indicate pockets of metastatic breast cancer. **h.** Metastatic load quantified as percentage of tumour to tissue area in vehicle control, and Eribulin-treated FCM-vac groups. Error bars represent SEM.

In contrast to whole-cell vaccines, the FCM-vac strategy eliminated the risk of residual tumour cell engraftment and allowed for a clean assessment of antitumour responses. Balb/cJRj mice were immunized twice with FCM-vac derived from either MYC^high^ or MYC-knockdown 4T1 cells **(Fig. 6d)**, then challenged with 1×10^5^ untreated 4T1 tumour cells injected subcutaneously into the right flank. Tumour growth was monitored every 2-3 days to evaluate vaccine efficacy.

In both MYC^high^ and MYC-knockdown vehicle-treated groups, 42.8% mice developed tumours exceeding the 400 mm^3^ threshold, whereas only 25% of mice vaccinated with FCM-vac derived from eribulin-treated cells surpassed this threshold **(Fig. 6e)**. Remarkably, none of the mice vaccinated with FCM-vac:Eri-shControl (MYC^high^, n=6) developed large tumours; one mouse remained completely tumour free, while the others showed slower tumour growth rates relative to other groups. In contrast, 50% of mice in the FCM-vac:Eri-shMYC (MYC knockdown) group developed tumours above the threshold. These findings closely recapitulate the MYC-selective effects observed in vitro and strongly demonstrate the tight correlation between MYC expression in cancer cells and immunogenicity of DAMPs released. Moreover, mice vaccinated with FCM-vac:Eri-shControl presented no metastatic nodules in the lungs, exhibiting the lowest metastatic burden among all groups tested **(Figs. 6f-h)**.

## DISCUSSION

Our study identifies microtubule inhibitors, including eribulin, as potent inducers of ICD in TNBCs, with a previously unappreciated dependence on MYC expression. Using a combination of in vitro DAMP reporter screens, DC activation assays, and ex vivo PDECs, we demonstrate that microtubule-targeting agents not only induce cytotoxicity preferentially in MYC^high^ cancer cells but also stimulate local immune activation, consistent with MYC-dependent synthetic lethal ICD. These findings build on prior work showing that ICD can be triggered by specific chemotherapeutics such as anthracyclines and oxaliplatin^59,60^, but expand the concept to microtubule-targeting agents in a MYC-dependent context.

While some standard chemotherapies are known to induce ICD, many were originally identified and explored for their antiproliferative effects rather than their capacity to stimulate immune responses^61,62^. Although biomarker profiling can predict tumour sensitivity to these agents, it often overlooks their immunomodulatory potential. Moreover, chemotherapy-associated toxicities, such as myelosuppression and systemic side effects^63^, highlight the clinical need for strategies that balance direct tumour cytotoxicity with immune activation^64,65^. In this context, microtubule inhibitors which selectively induce MYC-dependent ICD, provide both targeted cytotoxicity and local immune stimulation, thereby addressing a key limitation of conventional chemotherapeutics.

Mechanistically, microtubule inhibition triggered lytic cell death characterized by swelling, caspase-dependent GSDME cleavage, and release of immunostimulatory signals including HMGB1, CALR, and dsDNA, consistent with ICD such as pyroptosis^25,26^. This pattern explains the robust activation of DCs observed in co-culture assays, parallel to prior observations that DAMP release is essential for ICD-mediated adaptive immune responses^27^. Importantly, eribulin elicited similar cytokine signatures in PDECs, promoting pro-inflammatory and cytotoxic mediators while downregulating select immunosuppressive cytokines. In MYC^high^ PDECs, PGF and IL-22 were upregulated, whereas CSF2, IL-20, and TSLP were downregulated relative to MYC^low^ samples, reflecting a partial MYC-associated immunosuppressive effect, consistent with prior reports of MYC-driven immune evasion^18,19^. Despite these differences, the overall cytokine milieu indicated that eribulin-induced local immune activation remains intact even in the presence of MYC, though most differences were not statistically powered due to limited sample size.

In vivo, FCM vaccines derived from eribulin-treated endogenously MYC^high^ 4T1 cells conferred superior protection against tumour challenge compared with MYC-knockdown cells, directly demonstrating that MYC expression is essential for the release of immunogenic DAMPs during ICD. These findings underscore the dual role of MYC as both a driver of tumour proliferation and a determinant of susceptibility to microtubule-targeted ICD.

Overall, our work highlights the therapeutic potential of combining MYC-directed synthetic lethal strategies with ICD induction. Microtubule inhibitors, by selectively targeting MYC^high^ cells and promoting local immune activation, may serve as a foundation for next-generation chemotherapy regimens that integrate cytotoxicity with immune stimulation, addressing the limitations of conventional chemotherapeutics. Future studies should expand PDEC analyses to larger cohorts, explore combinatorial therapies integrating ICD-inducing microtubule inhibitors with immune checkpoint blockade, and investigate dosing regimens that maximize ICD while minimizing toxicity. Furthermore, elucidating the long-term immune memory induced by MYC-directed ICD may inform the clinical translation of immunotherapies for TNBC and other MYC-driven cancers.

## METHODS

### Image-based drug screen with FIMM Oncology collection library

MDA-MB-231 and HCC38 breast cancer cells were cultured in DMEM with 10% FBS, 2 mM L-glutamine, and 100 U/mL penicillin-streptomycin at 37°C with 5% CO₂. Lentiviral particles (Merck LentiBrite™ GFP-HMGB1 and RFP-CALR) were used to transduce cells sequentially at an MOI of 10 with 5 μg/mL polybrene. The GFP-HMGB1 transduced cells were cultured for 7 days, followed by CALR-RFP transduction. Double-positive cells were sorted using a SH800S Cell Sorter, ensuring 100% dual expression. Sorted cells were validated via EVOS Cell Imaging microscopy for HMGB1-GFP nuclear and CALR-RFP ER/plasma membrane localization. For the FIMM Oncology Collection drug screen, sorted cells were seeded in 384-well plates (1 × 10⁴ cells/well) for 24 hours, treated with library compounds for 48 hours, fixed with 4% paraformaldehyde, and stained with Hoechst 33342 (Thermo Scientific™ #62249) 1:10,000. Images were acquired using ImageXpress Pico Automated Cell Imaging System, with three fields per well selected as representative for analysis. Z score (z = (x – μ) / σ) was used to normalize the ICD-related parameters and select the top hits.

### Dendritic cell activation assay

CD14+ monocytes were isolated using positive magnetic separation according to manufacturer protocol (Miltenyi Biotec) and cultured in RPMI medium supplemented with 10% heat-inactivated FBS (Biowest), 100 U penicillin–streptomycin (Gibco), and 2mM L-Glutamine (Gibco) with added 100ng/mL Human granulocyte-macrophage colony-stimulating factor (GM-CSF) (Miltenyi Biotec), and 100 ng/mL Human IL-4 (Miltenyi Biotec) for 5 days, only adding media once in between. On day 5, DCs were harvested and 100μL of 1000 cells/μL plated on a 96-well flat-bottom plate. Samples were incubated with 100μL conditioned media from treated cells and DCs were harvested and analysed using flow cytometry on NovoCyte Quanteon after 24 hours. DCs expressing high levels of HLA-DR and CD86 were considered activated.

### Dendritic cell phagocytosis assay

Instead of the above, on day 5, 100 μL of 1000 cells/μL DCs were collected and co-cultured with CFSE stained TNBC cells pre-treated with drugs. DCs were harvested and analysed using flow cytometry on NovoCyte Quanteon after for 4-6 hours. CD45 markers were used to gate the DC populations from tumour cells.

### Cell death assays

HCC38-pInd20-MYC and Hs578T-pInd20-MYC were induced with 100nM doxycycline for 24 hours pre-treatment and then treated with drugs for another 24 hours. Cells were stained with Invitrogen SYTOX™ Green Dead Cell Stain (Invitrogen #S34860) 1:5000 and analysed on Novocyte Quanteon.

### Live-cell imaging

HCC38 ICD reporter cells were treated for 24 hours with 0.1 µM eribulin before live-cell imaging in an environmentally controlled chamber set at 37°C and 5% CO_2_. Imaging performed on Nikon Eclipse Ti microscope with Plan Apo λ 20x Ph2 DM objective, 0.75 aperture. Images captured at 1 per minute frequency for 5 hours.

### Patient Derived Explant Cultures

PDECs were generated with slight modifications to the methods previously described in Haikala et al, 2019. Fresh tissue was obtained from elective BC surgeries performed at the Helsinki University Central Hospital (Ethical permit: HUS/2697/2019, approved by the Helsinki University Hospital Ethical Committee). Patient participation in the study was authorized by signed informed consent. Collected tumour tissue was evenly portioned for formalin-fixed paraffin-embedded (FFPE) immunohistochemical analysis, quick-frozen at - 80°C for DNA/RNA/protein analysis, and processed for 3-dimensional culture. Explants were produced by incubating the samples overnight in collagenase A (3 mg/ mL; Sigma) and MammoCult Basal Medium (STEMCELL Technologies) with gentle shaking (130 rpm) at +37°C. MammoCult basal media was supplemented with MammoCult proliferation supplement (StemCell Technologies), 4 μg/ml heparin, 25 μg/mL gentamicin, 0.1 μg/ml Amphotericin B, and 100 U penicillin–streptomycin (Gibco). The resulting explants were collected via centrifugation at 353 rcf for 5 mins and washed once with 1× phosphate-buffered saline (PBS). Isolated explants were then embedded in Cultrex Reduced Growth Factor Basement Membrane Extract, Type 2 (R&D Systems) and plated on Nunc coated 8-well chamber slides (Thermo Scientific). Samples were cultured in MammoCult media as described above. PDECs generated were drug treated for 72 hours for cytokine profiling and flow cytometry assays.

### Immune biomarker and cytokine profiling

Olink Target 48 Cytokine and Olink Target 48 Immune Surveillance panels were combined to screen a total of 89 proteins involved in key immunological processes. 1µL conditioned medium from treated PDECs samples were used per panel. Refer to **Table S2.** for full list of markers screened.

### Generating concentrated fractionated conditioned medium vaccine (FCM-vac)

4T1-shMYC and 4T1-shControl mouse TNBC cell lines were cultured in RPMI 1640 media (Gibco) supplemented with 10% fetal bovine serum (FBS) (Gibco) 100 U penicillin– streptomycin (Gibco), and 2 mM L-Glutamine (Gibco). Cells were treated for 24 hours with either 1 μM eribulin, 1 μM vinorelbine, 1 μM paclitaxel, or 0.1% DMSO (Sigma) as vehicle control. After 24 hours, 15 mL conditioned medium was collected from treated cells and centrifuged at 1000g for 10 minutes to remove any live cells and cellular debris. The supernatant was then transferred to Amicon Ultra 15 mL centrifugal unit with 10 KDa molecular weight cutoff (MWCO) and centrifuged at 4000g for 1 hour. The supernatant was desalted with 1X PBS to remove traces of remaining drugs and salts in column eluent, resulting in 150 μL eluent containing 100X of concentrated factors present in initial conditioned medium. 10 KDa, 50KDa and 100KDa MWCO ultrafilters were used during experimental optimizations steps to investigate the most immunogenic fraction preceding final vaccine generation.

### In-vivo vaccination studies and tumour challenge

4-5 -week-old BALB/cJRj were obtained from Janvier Labs and housed in individual cages. Mice were acclimated between 1-2 weeks prior to the start of experimental procedures to minimize stress followed by ear-tagging. Mice were randomized and given a primary dose of 100 μL FCM-vaccine via subcutaneous injection into loose skin over the neck (n=7/ treatment group), followed by a booster dose of the vaccine after a 10-day interval. Mice were allowed to recover for 12 days before tumour challenge when 1 x 10^5^ cells/µL 4T1 cells suspended in 1X PBS were injected into loose skin over the right flank of each mouse. Tumour measurements were obtained using electronic callipers by two individual researchers who were not blinded to the treatment groups. Tumour volumes were calculated using formula V = 1/2 x L x W², where L = length and W = width. Mice were euthanized before any ethical limits were reached.

### Immunohistochemistry

Tissues and explant cultures were fixed with 4% paraformaldehyde (PFA) and embedded in paraffin. Samples were sectioned into 5 µm slices and deparaffinized. Heat-induced antigen retrieval was performed with a microwave oven in a citrate buffer solution, pH 6 (Dako). Endogenous peroxidase activity was blocked with 3% H_2_O_2_ for 20 minutes. Histochemical staining was carried out using standard techniques for IHC and anti-c-MYC antibody [Y-69] (Abcam #ab32072) at concentration of 1:300. Images were taken with a 3D HISTECH Panoramic 250 Flash III, with 20x (NA 0.8) air objective (Digital Microscopy and Molecular Pathology Unit, FIMM, Helsinki).

### Quantifying metastatic area in lungs

Two 5 µm tissue sections were cut 100 µm apart from FFPE lung blocks, slides were H&E stained and scanned as outlined above. Measurements taken as an average from regions of lung metastasis identified by two individual researchers, one of whom was blinded to experimental groups. Tumour burden was calculated as percentage of metastasis over total tissue area. Readings obtained using measurement tool within Aiforia Cloud version 6.2, build ID 6.2-680.

## ACKNOWLEDGEMENTS

We thank HiLIFE and Biocenter Finland funded Biomedicum Imaging Unit (BIU), Biomedicum Functional Genomics Unit (FuGU), Finnish Genome editing center (FINGEEC), Genome Biology Unit (GBU), Biomedicum Flow Cytometry Unit, FIMM Digital Microscopy and Molecular Pathology Unit, the Laboratory Animal Centre core facility at the University of and FIMM High Throughput Biomedicine Unit for their services. We thank Elina Hurskainen and Ellisiv Nyhamar for their technical support. The authors additionally thank Dr. Oliver Kepp and Prof. Guido Kroemer at Goustav Roussy for sharing the methodology behind ICD reporter cell line establishment. Schematic images were created with BioRender.com. Flow cytometry data was analysed on FlowJo v10 software. Graphs were generated and statistics calculated using GraphPad Prism 10. This work was funded by the Academy of Finland, Business Finland, the Finnish Cancer Organizations, the Sigrid Juselius Foundation, Jane and Aatos Erkko Foundation and RESCUER project, which has received funding from the European Union’s Horizon 2020 research and innovation programme under grant agreement No. 847912. This work was also supported by CDMRP W81XWH211-0773/-0774. Opinions, interpretations, conclusions, and recommendations are those of the author and are not necessarily endorsed by the Department of Defense. Support was received also from iCAN – Digital Precision Cancer Medicine Flagship (iCAN) and Finnish Red Cross Blood Service for funding this study. J.A. was funded by Doctoral Program in Biomedicine, The University of Helsinki Doctoral School.

## AUTHOR CONTRIBUTIONS

J.A, R.L, J.P and J.K. designed the concept of the study. J.A, R.L, A.H, L.I, M.S, M.P, R.T, N.S, J.M.A, B.P conducted the experiments. J.A, R.L, A.P, T.A.T, I.S, D.N analysed the data. M.M, J.M, P.E.K, L.N, T.M, contributed to clinical material collection and data analysis. J.A, R.L and J.K wrote the manuscript. A.G, P.M.M, J.P and J.K supervised the studies.

## STATISTICAL SIGNIFICANCE AND ETHICAL APPROVAL

Three to five biologically independent experiments were used in each study, and several assays were used for cross-validation. All animal experiments approved under National Animal Ethics Committee of Finland (License number: ESAVI/23309/2022) and mouse colonies maintained according to the protocols of the Experimental Animal Committee of the University of Helsinki. PDECs from live tumour samples obtained from elective breast cancer surgeries at the Helsinki University Central Hospital (Ethical permit: HUS/2697/2019 approved by the Helsinki University Hospital Ethical Committee) and taken with patient consent.

---

**Supplementary Figure 1.**
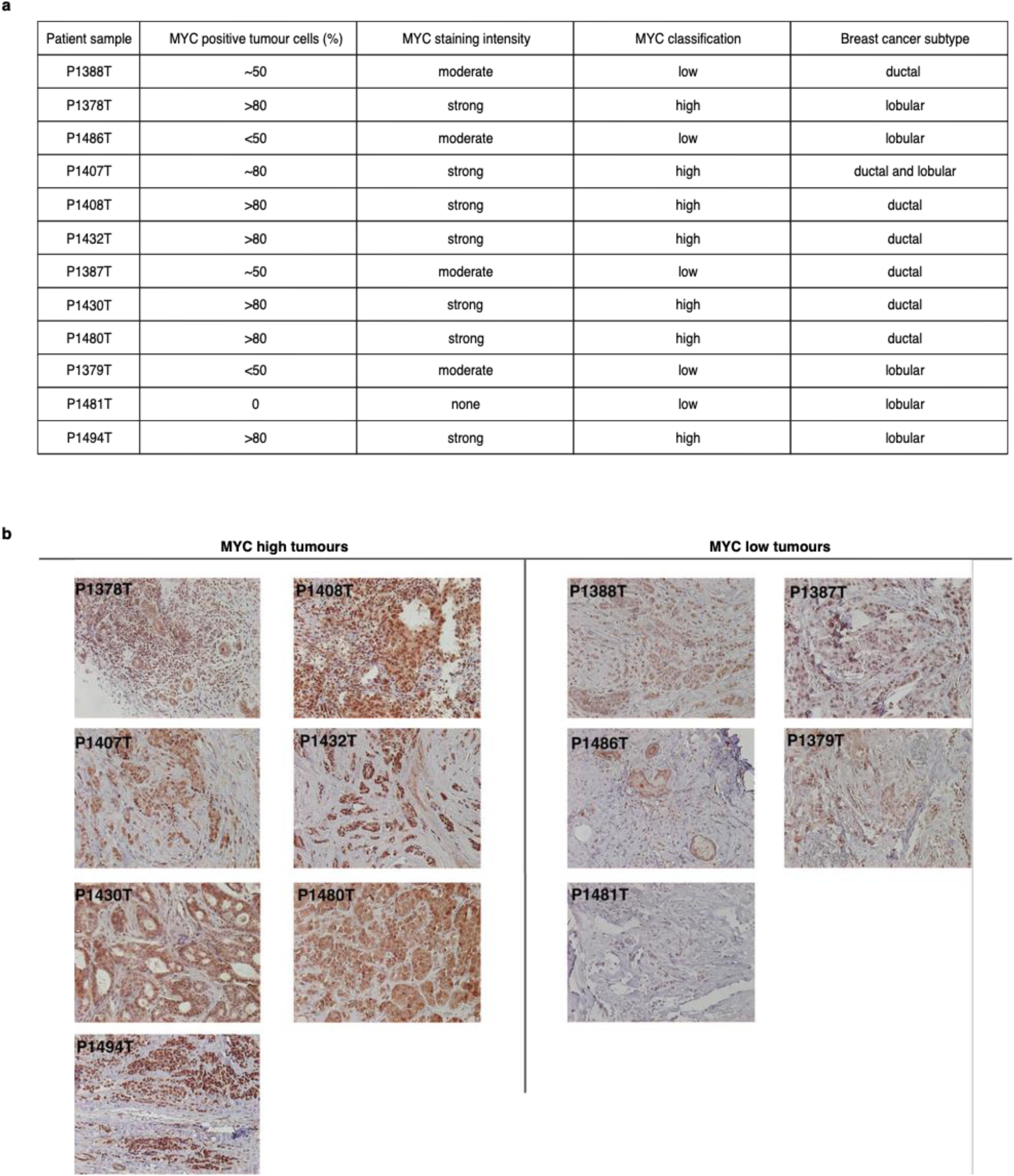
**a.** MYC classification of PDECs used in cytokine profiling. **b.** accompanying IHC stains of PDECs used in cytokine profiling.

**Supplementary Figure 2.**
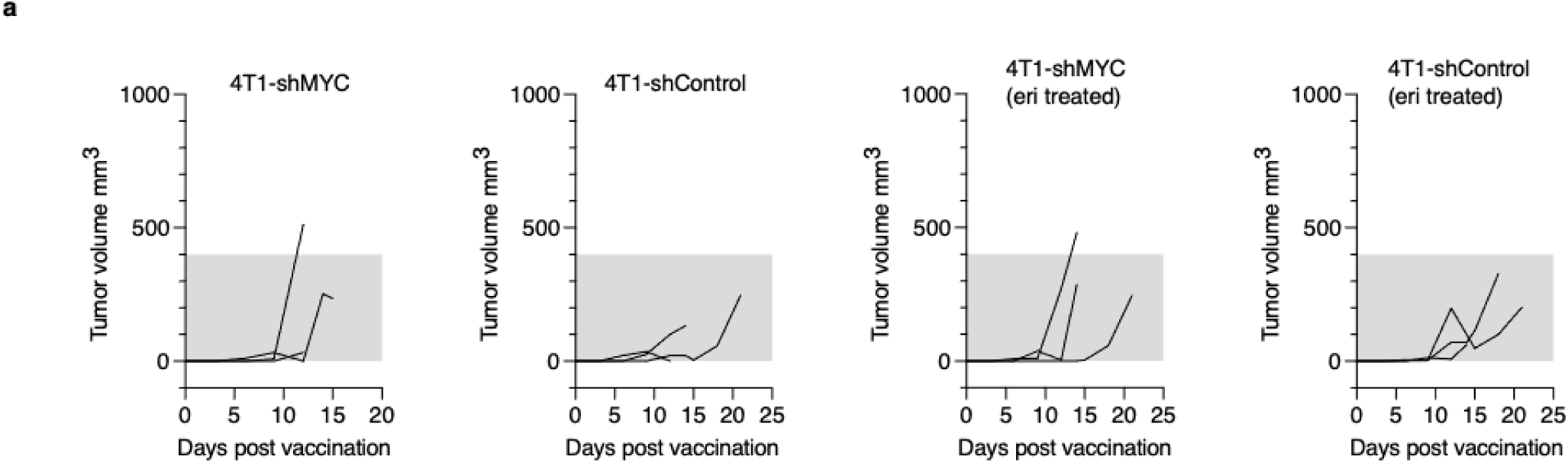
**a.** Graphs show tumour growth at vaccine site where 4T1 dying cell vaccine was given.

